# Fodder oats as catch crop: potential to reduce nitrogen losses from soil

**DOI:** 10.1101/2024.06.26.600925

**Authors:** Michael Kidson, Maria C. Hernandez-Soriano, Buhlebelive Mndzebele, Busiswa Ndaba, Rasheed Adeleke, Adornis D. Nciizah, Ashira Roopnarain

## Abstract

Reducing nitrogen (N) losses and associated nitrate (NO_3_^-^) leaching and nitrous oxide emissions from agricultural land is a critical target worldwide. This is particularly urgent in areas with low fertility soils and a climate that increases the risk of N loss, such as the arid and temperate regions of South Africa. Here, we assessed the potential of fodder oats (*Avena sativa*) as a winter catch crop to deplete residual N in a field laid fallow for the previous four years, where vetch had proliferated. The soil presented a high clay content (34-44%), with the main exchangeable bases being calcium and magnesium hence, ammonium (NH_4_^+^) deposited by the vetch was expected to be rapidly adsorbed and slowly released. A significant decrease in the concentrations of NO_3_^-^ (49%) and NH_4_^+^ (30%) throughout the soil profile (0-90 cm) was observed following harvest of the oats compared to the concentrations measured before sowing. The effectiveness of the oats to uptake both forms of N from top and deep soil layers enhances their potential to reduce N losses. Our results are useful to fill current knowledge gaps on N dynamics in understudied, vulnerable soils such as agricultural land in South Africa, and to advance crop rotation strategies that reduce risk of N leaching.

## 1. INTRODUCTION

Humanity is reliant on fertile agricultural soils to ensure food supply for the ever-growing population, but at least 33% of all croplands are moderately or highly degraded globally (Itps, 2015; Davis et al., 2023). This threat to food security is particularly severe in countries like South Africa (Ighodaro et al., 2016; Daniell and van Tonder, 2023; Roopnarain et al., 2024). Mitigation of low soil fertility through application of synthetic nitrogen (N) fertilizers currently supports the production of half of the food humans consume (Ritchie, 2017), but half of the N-fertilizer inputs are lost to the environment worldwide (Havlin, 2020; Duncombe, 2021) with a detrimental impact on yields, agricultural wealth and the environment. Nitrate (NO_3_^-^) leaching contributes to eutrophication and the emission of nitrous oxide plays a major role in global warming (Rezaei Rashti et al., 2015; Henryson et al., 2020; Maaz et al., 2021). Nitrogen leaching from crop land in South Africa is particularly acute during high rainfall events (Tongwane et al., 2020). Consequently, mitigating N losses from agricultural soils, particularly through leaching of nitrates to deeper soil layers and groundwater, is a critical economic and environmental demand.

Efforts to improve N use efficiency (NUE) to enhance crop production and minimise environmental pollution have been proven challenging (Timilsena et al., 2015; Tian et al., 2021). Strategies such as use of nitrification inhibitors and organic amendments have been reported as rather ineffective in mitigating N-fertilizer losses (Coskun et al., 2017; Bossolani et al., 2023). Nitrogen recovery by catch crops in soils with high N residual content is an increasingly recognized strategy to optimize NUE and minimize N losses (Abdalla et al., 2019; Malcolm et al., 2022). Catch crops can utilize excess soil nutrients and gradually release them to the soil when decomposing during the following cropping season (Berntsen et al., 2006; Constantin et al., 2011). Besides their nutritional advantages (Rasane et al., 2015), oats (*Avena sativa*) are a reportedly effective catch crop candidate (Gentsch et al., 2022) to decrease NO_3_^-^ leaching and enhance soil health and crop profitability (Talanow et al., 2021; Vogeler et al., 2023), particularly relevant in semiarid regions like South Africa (Milton and Dean, 1995; Acharya et al., 2022). Fodder oats are highly valuable for farming systems, especially for smallholders (FAO, 2004). Additionally, oats as a catch crop provides protection against soil borne diseases (Leveau et al., 2019) that affect cash crops like wheat (Tadesse et al., 2019; Shew et al., 2020). The present study therefore aimed to investigate the potential of fodder oats as a winter catch crop on a South African soil.

Previous studies have focused on the impact of catch crops on N levels in the topsoil, with limited information on their ability to access and take up N from deeper soil horizons. Here, we assess the impact of fodder oats through the soil profile. Also, this study focused on a soil with high clay content (34-44%), prevalent in South Africa (Sumner, 2015; Daniell and van Tonder, 2023), where ammonium (NH_4_^+^) is expected to be rapidly adsorbed onto clay minerals and slowly released over time (Yu et al., 2023).

## 2. MATERIALS AND METHODS

A field study was conducted in 2022 in South Africa (Agricultural Research Council research farm, Brits (25,5849 N, 27,7692 E), to assess the ability of fodder oats to reduce N levels in the soil profile. The field was laid fallow for four years prior to this study, with mostly couch grass (*Cynodon dactylon*) and broad-leafed purple vetch (*Vicia sativa*) growing in the field.

Fodder oats were obtained from a local provider and sown in May 2022. No N-fertilizer or herbicides were applied throughout the study. The oats were irrigated using sprinklers with water being sourced from the Hartbeespoort dam, with a recorded low N content. Three sets of soil samples were collected throughout the study using a Dutch soil auger. Samples were collected at three depths (0-30 cm, 30-60 cm, and 60-90 cm) at relevant times: prior to sowing (May 2022), at plant ripening (September 2022) and following harvest (October 2022). Soil fertility and soil texture were determined from the first set of samples through chemical characterization (Jackson, 2005; Hailu et al., 2015) and particle size analysis (Murano et al., 2015). Nitrate and NH_4_^+^ content in the soil profile were determined for the first and third sets of samples by soil extraction with KCl 1 M and measured in a SEAL Analytical AAIII segmented flow, colorimetric, auto-analyser (Technicon AutoAnalyzer, n.d.). To determine the impact of the oats on soil microbial activity, N potential release (NPR) and potentially mineralisable N (PMN) (Moore et al., 2019) were determined using the Solvita CO_2_ burst test (Haney et al., 2008; Chahal et al., 2021) for the second and third set of samples.

## 3. RESULTS AND DISCUSSION

The soil was classified as Cambic calcisol (Chromic) (van Huyssteen, 2020), Kimberley form (Classification Working Group, n.d.; van Zijl et al., 2020), according to the particle size analysis (Table S1), this being consistent with the arid climate of the area (Köppen, 1931; Cui et al., 2021). Base saturation analysis indicated that for this soil exchangeable sites are occupied mainly by calcium (Ca) and magnesium (Mg) (Table 1). The ratio of potassium (K) to Mg suggests a Mg-induced K deficiency (Hailu et al., 2015). The ratio Ca to Mg and the Na saturation lower than 3% suggest that the soil has potential for self-mulching (Laker and Nortjé, 2019) and a moderate tendency to disperse (Rengasamy et al., 1986). The clay content ranged 34-44% through the soil profile (Table S1), with a gradual decrease of resistivity with depth (Table 1) likely related to the clay content increase (Long et al., 2012). The low soil resistivity values recorded can be partly attributed to the proliferation of vetch over the four years before the study (Gabriel et al., 2021).

**Table 1.** Exchangeable bases, cation exchange capacity (CEC), base saturation percentage, K to Mg ratio, Ca to Mg, available phosphorus (P), resistivity (R) and pH of the study site at three depths (n=3).

| Depth<br>cm | P<br>mg kg<sup>-1</sup> | Exchangeable bases cmol<sub>(+)</sub> kg<sup>-1</sup> |  |  |  | K:Mg | Ca:Mg | CEC<br>cmol<sub>(+)</sub> kg<sup>-1</sup> | Base saturation % CEC |  |  |  | R<br>ohm.cm | pH* |
| --- | --- | --- | --- | --- | --- | --- | --- | --- | --- | --- | --- | --- | --- | --- |
|  |  | Ca | Mg | K | Na |  |  |  | Ca | Mg | K | Na |  |  |
| 0-30 | 3.5 | 1.3 | 9.5 | 0.30 | 0.36 | 0.03 :1 | 0.14 :1 | 11.5 | 11 | 83 | 2.6 | 3.2 | 510 | 7.23 |
|  | 2.8 | 12 | 7.2 | 0.24 | 0.38 | 0.03 :1 | 1.67 :1 | 19.8 | 61 | 36 | 1.2 | 1.9 | 640 | 7.33 |
|  | 13 | 12 | 7.9 | 0.35 | 0.35 | 0.04 :1 | 1.53 :1 | 20.6 | 58 | 38 | 1.7 | 1.7 | 720 | 7.13 |
|  | 11 | 12 | 6.8 | 0.33 | 0.32 | 0.05 :1 | 1.76 :1 | 19.5 | 62 | 35 | 1.7 | 1.6 | 560 | 7.3 |
| 30-60 | 0.13 | 18 | 12.0 | 0.32 | 0.71 | 0.03 :1 | 1.50 :1 | 31.0 | 58 | 39 | 1.0 | 2.3 | 340 | 7.86 |
|  | 0.58 | 19 | 8.3 | 0.30 | 0.57 | 0.04 :1 | 2.28 :1 | 28.2 | 67 | 30 | 1.1 | 2.0 | 530 | 7.46 |
|  | 1.1 | 13 | 9.6 | 0.26 | 0.53 | 0.03 :1 | 1.36 :1 | 23.4 | 56 | 41 | 1.1 | 2.3 | 650 | 7.54 |
|  | 0.42 | 15 | 9.8 | 0.28 | 0.50 | 0.03 :1 | 1.53 :1 | 25.6 | 59 | 38 | 1.1 | 2.0 | 480 | 7.54 |
| 60-90 | 0.05 | 30 | 13.0 | 0.35 | 0.81 | 0.03 :1 | 2.31 :1 | 44.2 | 68 | 29 | 0.8 | 1.8 | 360 | 8.22 |
|  | 0.08 | 14 | 7.9 | 0.26 | 0.50 | 0.03 :1 | 1.78 :1 | 22.6 | 62 | 35 | 1.1 | 2.2 | 430 | 7.86 |
|  | 0.30 | 12 | 9.7 | 0.26 | 0.54 | 0.03 :1 | 1.24 :1 | 22.5 | 53 | 43 | 1.1 | 2.4 | 630 | 7.69 |
|  | 0.18 | 19 | 11.3 | 0.32 | 0.62 | 0.03 :1 | 1.68 :1 | 31.3 | 61 | 36 | 1.0 | 2.0 | 410 | 7.86 |

The initial NO_3_^-^ concentrations in the soil studied (Figure 1) were comparable to those reported by previous studies in NE South Africa (Ntalo et al., 2022), while the NH_4_^+^ concentrations were above average, likely due to rhizodeposition of N by the vetch (Ozpinar and Baytekin, 2006) and the adsorption of N on the clay fraction. Soils in the area are rich in smectite, which hold high quantities of essential nutrients that are easily available for plant uptake (Daniell and van Tonder, 2023). This is confirmed by the high percentage of exchangeable Ca^2+^ and Mg^2+^ (Table 1) which reportedly increase the soil capacity to adsorb N (Dontsova et al., 2005; Nieder et al., 2011).

**Figure 1.**
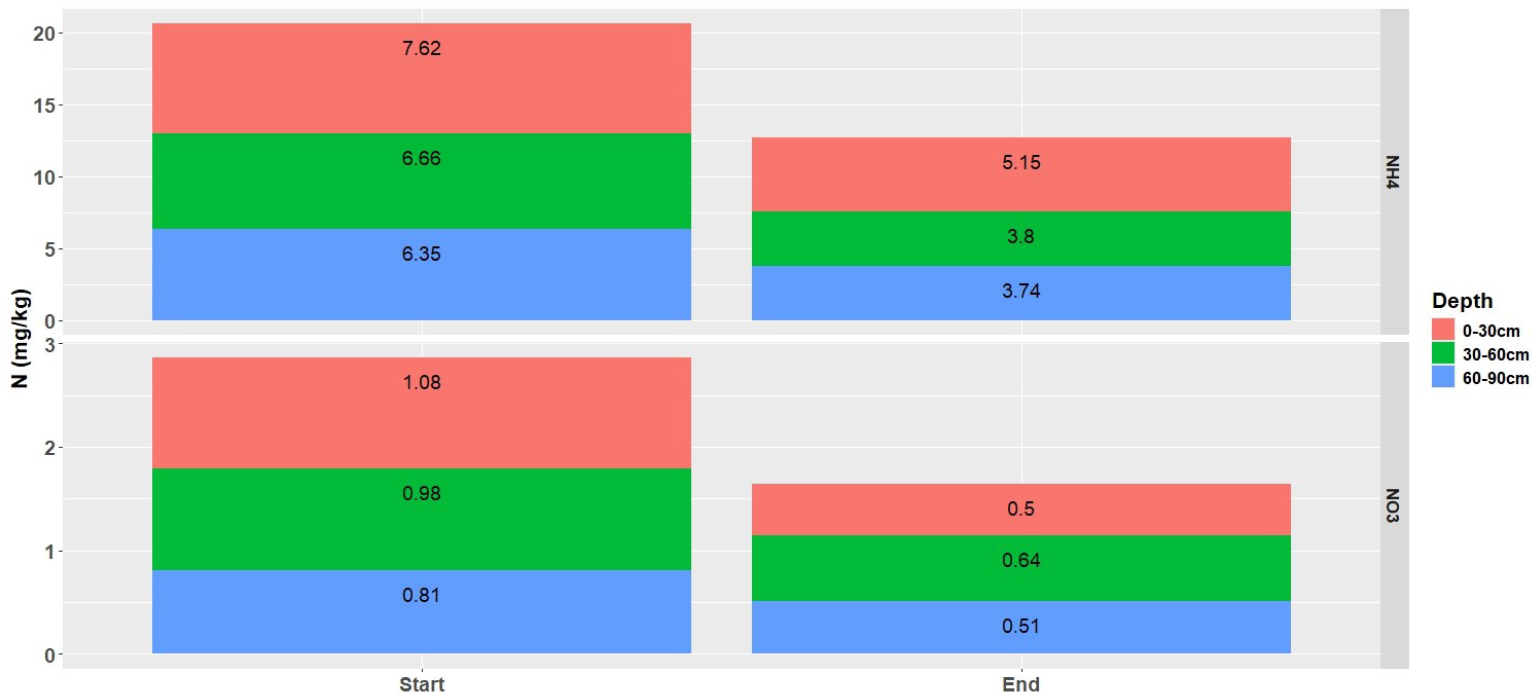
Average values (n=3) of ammonium (NH_4_^+^) and nitrate (NO_3_^-^) across the soil profile before (Start) and after (End) growing oats. Significant decrease was confirmed for both forms of nitrogen (Bonferroni t-test, p=2.31e-5 for (n=3) and p=4.3e-4 for NO_3_^-^), for the three depths sampled.

Sowing fodder oats over winter significantly decreased the concentrations of NO_3_^-^ (49%) and NH_4_^+^ (30%) at the three depths of the soil profile studied (Figure 1) (Bonferroni t-test, p=2.31e-5 for NH_4_^+^ and p=4.3e-4 for NO_3_^-^). These results are consistent with previous research reporting that oats can access N from deeper soil levels than other crops (Af Geijersstam and Mårtensson, 2006) and with previous studies showing a reduction of N losses by planting oats over winter (Michel et al., 2021; Malcolm et al., 2022), therefore confirming their potential to reduce N leaching (Carey et al., 2018). No significant differences were obtained for the NH_4_^+^/NO_3_^-^ ratio (Figure 2), with only a minor increase of the ratio in the top layer, which indicates that oats take up both forms of N (van Lierop and Tran, 1980). The uptake of NH_4_^+^ adsorbed to the clay minerals by the oats can be explained by different mechanisms. Firstly, the decrease of NH_4_^+^ ions in the soil solution might lead to their diffusion from the clay minerals but the dynamics remain unclear (Nieder et al., 2011). Second, recently adsorbed NH_4_^+^ such as that likely deposited by the vetch can be released over the three following months and therefore it would have become available for the oats to uptake (Kowalenko and Cameron, 1976; Scherer, 1993). Thirdly, the roots of the oats likely released exudates that might have promoted the desorption and depletion of NH_4_^+^ by the plant (Trofymow et al., 1987; Fisk et al., 2015). Release of exudates is also expected to affect the soil biological activity in relation to N availability (Zhalnina et al., 2018; Morales et al., 2023). Although the results from the Solvita CO_2_ burst test were not conclusive due to the variability of the measurements (Table 2, Table S2), we determined a significant decrease in NH_4_^+^ as plants grew to full maturity, which confirms that the NH_4_^+^ released by desorption was the main source of N for the oats. In summary, growing oats through winter can significantly decrease the risk of N losses by leaching or gas emission, particularly when grown in land previously laid fallow (Tongwane et al., 2020).

**Table 2.** Average soil microbial activity (n=3) in the topsoil (0-30 cm) determined by the Solvita CO_2_ burst test and Potentially Mineralisable Nitrogen (PMN) analysis. Soil microbial activity and nitrogen potential release (NPR) were estimated according to the scale of microbial soil activity (Solvita® Guidelines, 2013) and consistently medium-high at the three sampling times.

| Sampling date | CO<sub>2</sub> (ppm) | NH<sub>4</sub>-N (mg/kg) | NO<sub>3</sub>-N (mg/kg) | PMN (mg/kg) |
| --- | --- | --- | --- | --- |
| Sep-2022 | 64 ± 24<sup>a</sup> | 1.57 ± 0.21<sup>a</sup> | 1.19 ± 0.54<sup>a</sup> | 1.05 ± 1.17<sup>a</sup> |
| Oct-2022 | 77 ± 28<sup>a</sup> | 1.20 ± 0.14<sup>b</sup> | 0.64 ± 0.10<sup>a</sup> | 1.37 ± 1.34<sup>a</sup> |

**Figure 2.**
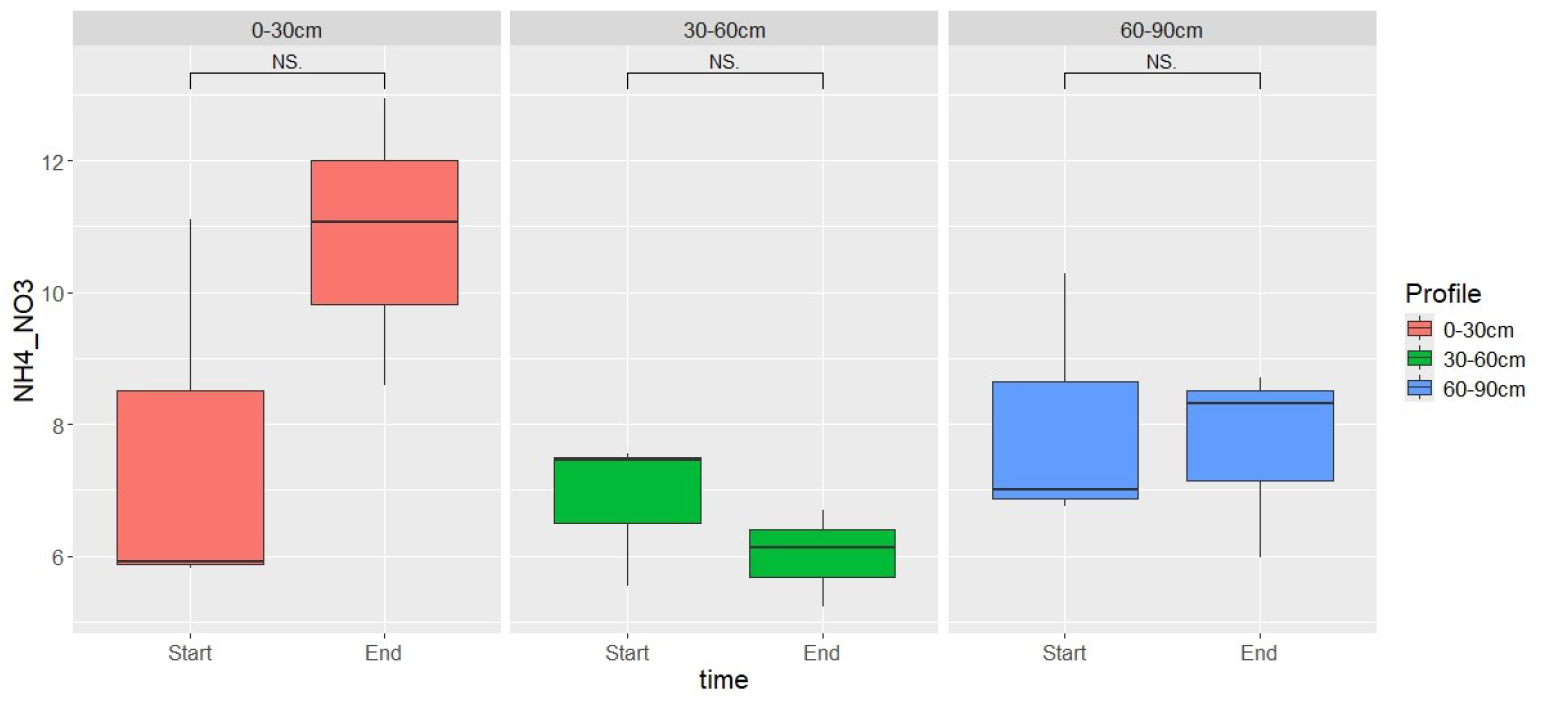
Ammonia (NH_4_^+^) to nitrate (NO_3_^-^) ratios (n=3) across the soil profile before (Start) and after (End) growing oats. No significant differences were observed but significant decrease was confirmed for both forms of nitrogen (t-test, p>0.05)

## 4. CONCLUSION

Fodder oats can be used as an effective catch crop over winter to deplete N from soil since this crop can uptake both NO_3_^-^ and NH_4_^+^ recently adsorbed on to the clay minerals. This makes them particularly suitable for soils with high clay content where NH_4_^+^ is rapidly adsorbed and slowly released. Our results confirm that oats can uptake both forms of N from deep soil layers, which enhances their potential to reduce N losses by leaching. The results presented are useful to fill current knowledge gaps on N dynamics in understudied, low fertility soils such as agricultural land in South Africa, and to develop crop rotation strategies that reduce risk of N leaching.

## AUTHOR CONTRIBUTIONS

**Michael Kidson**: Conceptualisation, writing, editing, data analysis. **Maria C. Hernandez-Soriano:** writing - review and editing; data analysis. **Buhlebelive Mndzebele:** sampling, analysis, review. **Busiswa Ndaba:** review. **Rasheed Adeleke:** review. **Adornis Nciizah:** review & editing. **Ashira Roopnarain**: review, editing.

## ACKNOWLEDGEMENTS

The project is funded by the European Joint Programme Soil ERA-NET (HORIZON 2020) and the New Zealand Government to support the objectives of the Global Research Alliance on Agricultural Greenhouse Gases (Project P07000201). MCHS gratefully acknowledges the support of the Biotechnology and Biological Sciences Research Council (BBSRC); her research is funded by the BBSRC project BB/X003000/1.

## DATA AVAILABILITY STATEMENT

The data that support the findings of this study are partially available from the supporting material and further from the corresponding author upon reasonable request.

