## Supplemental Tables 1-2 for "Fodder oats as catch crop: potential to reduce nitrogen losses from soil"

**Contents**

**Table 1**. Particle size analysis. Soil was classified as Kimberley soil (Classification Working Group, n.d.) and Cambic calcisol (Chromic) (van Huyssteen 2020). .

**Table 2**. Soil microbial activity and nitrogen potential release (NPR) determined by the Solvita $CO_2$ burst test and Potentially Mineralisable Nitrogen (PMN) analysis.

**Table S1**. Particle size analysis. Soil was classified as Kimberley soil (Classification Working Group, n.d.) and Cambic calcisol (Chromic) (van Huyssteen 2020)**.**

| Depth cm | Sand % | Silt % | Clay % |
|---|---|---|---|
| 0-30 | 58 | 4 | 38 |
| | 64 | 2 | 34 |
| | 60 | 6 | 34 |
| | 60 | 6 | 34 |
| 30-60 | 54 | 4 | 42 |
| | 58 | 6 | 36 |
| | 54 | 6 | 40 |
| | 56 | 6 | 38 |
| 6-90 | 50 | 8 | 42 |
| | 54 | 8 | 38 |
| | 56 | 8 | 36 |
| | 48 | 8 | 44 |

**Table S2**. Average soil microbial activity (n=3) in the topsoil (0-30 cm) determined by the Solvita CO2 burst test and Potentially Mineralisable Nitrogen (PMN) analysis. Soil microbial activity and nitrogen potential release (NPR) were estimated according to the scale of microbial soil activity (Solvita® Guidelines, 2013) and consistently medium-high at the three sampling times.

| Depth cm | $CO_2$ ppm | Soil microbial activity | NPR* | $NH_4$-N mg $kg^{-1}$ | $NO_3$-N mg $kg^{-1}$ | PMN mg $kg^{-1}$ |
|---|---|---|---|---|---|---|
| | 59.7 | Medium | Some N-min | 1.35 | 1.34 | 0.12 |
| Sep-2022 | 43.5 | High | Strong N-min | 1.57 | 0.59 | 0.65 |
| | 90.2 | Medium | Some N-min | 1.78 | 1.65 | 2.37 |
| | 108 | Medium | Some N-min | 1.04 | 0.63 | 0.14 |
| Oct-2022 | 54.5 | Medium | Some N-min | 1.26 | 0.54 | 1.18 |
| | 68.7 | High | Strong N-min | 1.30 | 0.75 | 2.80 |

*According to the scale of microbial soil activity (Solvita® Guidelines, 2013) “some N-min” is equivalent to 11-22 kg $ha^{-1}$ and “strong N-min” is equivalent to 84-112 kg $ha^{-1}$.
